# Splicing Factor SF3B2 Regulates Cardiomyocyte Calcium Handling Through Alternative Splicing of Cardiac Ion Channel Genes

**DOI:** 10.64898/2026.07.25.740735

**Authors:** Sean Murphy, Hanwen Wang, Nadine Zureick, Misato Koakutsu, David Suh, Dong Ik Lee, Chulan Kwon

## Abstract

**Background:** Alternative splicing is a critical determinant of protein diversity in the heart, where it drives the postnatal functional maturation of cardiomyocytes and specifies the ion-channel and calcium-handling isoforms required for mature contractile function; dysregulated splicing programs have in turn been implicated in cardiomyopathies and arrhythmias. However, the splicing regulators that control cardiomyocyte calcium handling remain largely unknown.

**Objective:** We systematically screened 276 splicing factor genes in human induced pluripotent stem cell-derived cardiomyocytes (hiPSC-CMs) to identify regulators of calcium handling, and we selected SF3B2, a core U2 snRNP spliceosome component, for detailed follow-up based on complex-level enrichment and prior identification by Murphy et al. 2021 [1].

**Methods:** A high-throughput siRNA screen targeting 276 splicing factor genes was performed in hiPSC-CMs using 384-well calcium transient imaging. SF3B2 knockdown was followed by deep bulk RNA-seq analysis (n = 3 per group) for differential gene expression with DESeq2 and alternative splicing quantification with rMATS. RNA immunoprecipitation sequencing (RIP-seq) using an epitope-tagged SF3B2 construct was performed to identify direct mRNA binding targets of SF3B2.

**Results:** The screen identified multiple U2 snRNP components, including SF3A2, SF3B1, and SF3B4—as regulators of calcium transient duration. SF3B2 was previously identified as a regulator of contractility in this screen [1], and the enrichment of its complex partners above the significance threshold in the current CTD75 analysis supported its selection for follow-up characterization. SF3B2 knockdown resulted in 2,683 differentially expressed genes (adjusted *p* < 0.05), with downregulated genes enriched in cell cycle and DNA replication pathways. Alternative splicing analysis revealed significant changes across all five rMATS event types in 36 genes within the cardiac muscle cell action potential involved in contraction gene ontology term (GO:0086002), including calcium channel (*CACNA1C*, *CACNA1D*, *CACNA2D1*), potassium channel (*KCNH2*, *KCNQ1*), and sodium channel (*SCN5A*) genes, although the large number of affected genes is consistent with broad spliceosomal perturbation and a formal enrichment test would be needed to determine whether cardiac action potential genes are preferentially affected. Integration of RIP-seq data (597 SF3B2-enriched transcripts, FDR < 0.05) with splicing analysis identified *CACNA2D1,* which encodes an auxiliary subunit of L-type calcium channels, as both directly bound by SF3B2 and alternatively spliced upon knockdown.

**Conclusion:** These findings identify SF3B2 as a regulator of cardiomyocyte calcium handling and suggest that SF3B2-dependent missplicing of *CACNA2D1* may link core spliceosome function to the splicing programs underlying cardiomyocyte functional maturation.

**Significance Statement:** This study provides the first systematic functional screen of splicing factors in cardiomyocyte calcium handling and identifies SF3B2, a U2 snRNP subunit, as a regulator of cardiac ion channel splicing. Transcriptomic, splicing, and RNA binding data converge on *CACNA2D1*, identifying a route by which a core splicing factor shapes cardiac contractility, extending cardiac splicing regulation beyond the accessory RBPs studied to date.

## Introduction

Most multi-exon human genes produce multiple transcript isoforms through alternative pre-mRNA splicing [2], and in the heart, tightly regulated splicing programs specify the functionally distinct isoforms of sarcomeric proteins, ion channels, and calcium handling machinery that underlie developmental transitions and adult contractile function [3, 4]. These splicing transitions are integral to the postnatal functional maturation of cardiomyocytes, and organ-wide splicing switches have recently been shown to drive maturation of the postnatal heart [5]. This program is incompletely recapitulated in vitro: human induced pluripotent stem cell-derived cardiomyocytes (hiPSC-CMs) remain stalled at a fetal-like state, with immature calcium handling, sarcomeric organization, and electrophysiology that limit their utility for disease modeling and regenerative applications [6, 7, 8]. Trajectory analyses indicate that hiPSC-CMs fail to execute the perinatal maturation program engaged by cardiomyocytes in vivo [8], and multiple regulatory inputs, including hormonal and neuronal cues [9] and RNA-splicing regulators such as RBFox1 [10] are abnormally regulated or absent under standard culture conditions. Dysregulation of alternative splicing has also been linked to dilated cardiomyopathy, hypertrophic cardiomyopathy, and cardiac arrhythmias [11, 12]. Pre-mRNA splicing is catalyzed by the spliceosome, a dynamic multi-megadalton ribonucleoprotein complex composed of five small nuclear ribonucleoprotein particles (U1, U2, U4, U5, and U6 snRNPs) and numerous associated factors [13]. How individual spliceosome components contribute to cardiomyocyte-specific splicing programs is unknown.

Most cardiac splicing studies to date have examined accessory RNA-binding proteins (RBPs). RBM20 mutations cause dilated cardiomyopathy through missplicing of titin and other sarcomeric transcripts [11]. MBNL1 regulates fetal-to-adult splicing transitions in the heart [3], RBFOX2 controls splicing of cytoskeletal and ion channel transcripts [12], and QKI was recently identified as a critical regulator of cardiac myofibrillogenesis through pre-mRNA alternative splicing [14]. These studies have established that individual splicing regulators can exert gene-specific control over cardiac transcripts. Whether core spliceosome components can selectively regulate splicing of functionally related gene sets, such as those controlling calcium handling, which driveds cardiomyocyte contractility through L-type calcium channels, ryanodine receptors, SERCA2a, and associated regulatory proteins [15], is less investigated.

To address this question, we performed a high-throughput siRNA screen targeting 276 splicing factor genes in human induced pluripotent stem cell-derived cardiomyocytes (hiPSC-CMs), using calcium transient duration (CTD) as a functional readout. SF3B2 was identified as a regulator of cardiomyocyte contractility in this screen, as reported in Murphy et al. 2021 [1]. In the current CTD75 analysis, multiple U2 snRNP components that are SF3B2 complex partners—including SF3A2, SF3B1, and SF3B4—scored above the significance threshold. Based on this complex-level enrichment and SF3B2’s prior identification, we selected SF3B2 for follow-up characterization. SF3B2 is one of seven subunits of the SF3b subcomplex of the U2 snRNP, which recognizes the branch point sequence during the earliest steps of spliceosome assembly [16, 17]. Here, we characterized the transcriptomic and splicing consequences of SF3B2 knock down in hiPSC-CMs and used RIP-seq to identify its direct mRNA targets.

## Materials and Methods

### hiPSC-CM Differentiation and Culture

Human induced pluripotent stem cells (WTC11 line) were differentiated into cardiomyocytes using a small-molecule Wnt modulation protocol as previously described [18]. Cardiomyocytes were cultured in RPMI 1640 supplemented with B27 and used for experiments at day 25 post-differentiation. Spontaneous beating was confirmed visually prior to experiments.

### High-Throughput Calcium Imaging Screen

An siRNA library targeting 276 splicing factor genes (Dharmacon) was arrayed in 384-well plates. Day 25 hiPSC-CMs were replated into library plates, transfected, and incubated for 72 hours. Calcium transient duration at 75% repolarization (CTD75) was measured using live calcium imaging. CTD75 values were normalized to the median of on-plate scrambled siRNA controls. Hits were identified based on statistical significance relative to plate controls. Screen replicates and quality metrics (including Z’-factor) are detailed in Murphy et al. 2021 [1]. The screen design and validation of SF3B2 as a hit have been described previously [1].

### SF3B2 Knockdown and RNA-seq

SF3B2 knockdown was performed using Dharmacon ON-TARGETplus SMARTpool siRNA and Lipofectamine RNAiMAX in day 25 hiPSC-CMs. Knockdown efficiency was validated by quantitative real-time PCR (qPCR), confirming 78% +/- 6% (mean +/- SD, n = 3) reduction of SF3B2 mRNA relative to scrambled siRNA controls. Cells were harvested 48 hours post-transfection. Total RNA was extracted from n = 3 biological replicates per condition and sequenced by Novogene (approximately 50 million paired-end stranded reads per sample). Reads were aligned to the human reference genome (GRCh38) using HISAT2, quantified with featureCounts, and analyzed for differential gene expression using DESeq2 with batch correction [19]. Alternative splicing was quantified using rMATS (replicate Multivariate Analysis of Transcript Splicing), which evaluates five event types: skipped exon (SE), retained intron (RI), alternative 3’ splice site (A3SS), alternative 5’ splice site (A5SS), and mutually exclusive exons (MXE) [20]. Significant splicing events were defined as FDR < 0.05 and |deltaPSI| > 0.1. This threshold is commonly used in the field to filter statistically significant but quantitatively minor splicing changes, ensuring that reported events reflect biologically meaningful isoform shifts.

### RNA Immunoprecipitation Sequencing (RIP-seq)

hiPSC-CMs were transduced with AAV6 expressing either CMV-SF3B2-3xHA or CMV-GFP (control). Cells were harvested 72 hours post-transduction and RNA immunoprecipitation was performed using the Magna RIP kit (Millipore) with anti-HA antibody (n = 3 biological replicates per condition). Immunoprecipitated RNA was sequenced by Novogene and analyzed using DESeq2 to identify transcripts significantly enriched in the SF3B2-3xHA pulldown relative to GFP control (FDR < 0.05).

### Statistical Analysis

All statistical analyses were performed using R/Bioconductor. Differential gene expression was assessed using DESeq2 (adjusted *p* < 0.05). Alternative splicing significance was determined by rMATS (FDR < 0.05). RIP-seq enrichment was assessed by DESeq2 (FDR < 0.05). Gene ontology enrichment analysis was performed using PantherGO [21].

## Results

### High-Throughput Splicing Factor Screen Identifies U2 snRNP Components as Regulators of Cardiomyocyte Calcium Handling

To systematically identify splicing factors required for cardiomyocyte calcium handling, we performed an siRNA screen targeting 276 splicing factor genes in hiPSC-CMs, using calcium transient duration at 75% repolarization (CTD75) as a functional readout (Fig. 1A). Through this screen, we identified multiple splicing factors that significantly altered the duration of the calcium transient when their expression was suppressed (Fig. 1B). Among the top hits were the RNA-binding proteins QKI, LSM2, and SF3A2. Several components of the U2 snRNP spliceosome complex, including SF3A2, SF3B1, and SF3B4, scored above the significance threshold. SF3B2 had previously been identified as a regulator of cardiomyocyte contractility [1]; the enrichment of multiple SF3B2 complex partners above the CTD75 significance threshold in the present analysis supported its selection for detailed follow-up. Mapping the screen hits onto the cryo-EM structure of the human 17S U2 snRNP complex [16] revealed that the identified subunits clustered within the SF3a and SF3b subcomplexes, with SF3A2, SF3A3, SF3B1, SF3B2, SF3B4, and SF3B5 all represented among the screen hits (Fig. 1C).

**Figure 1.**
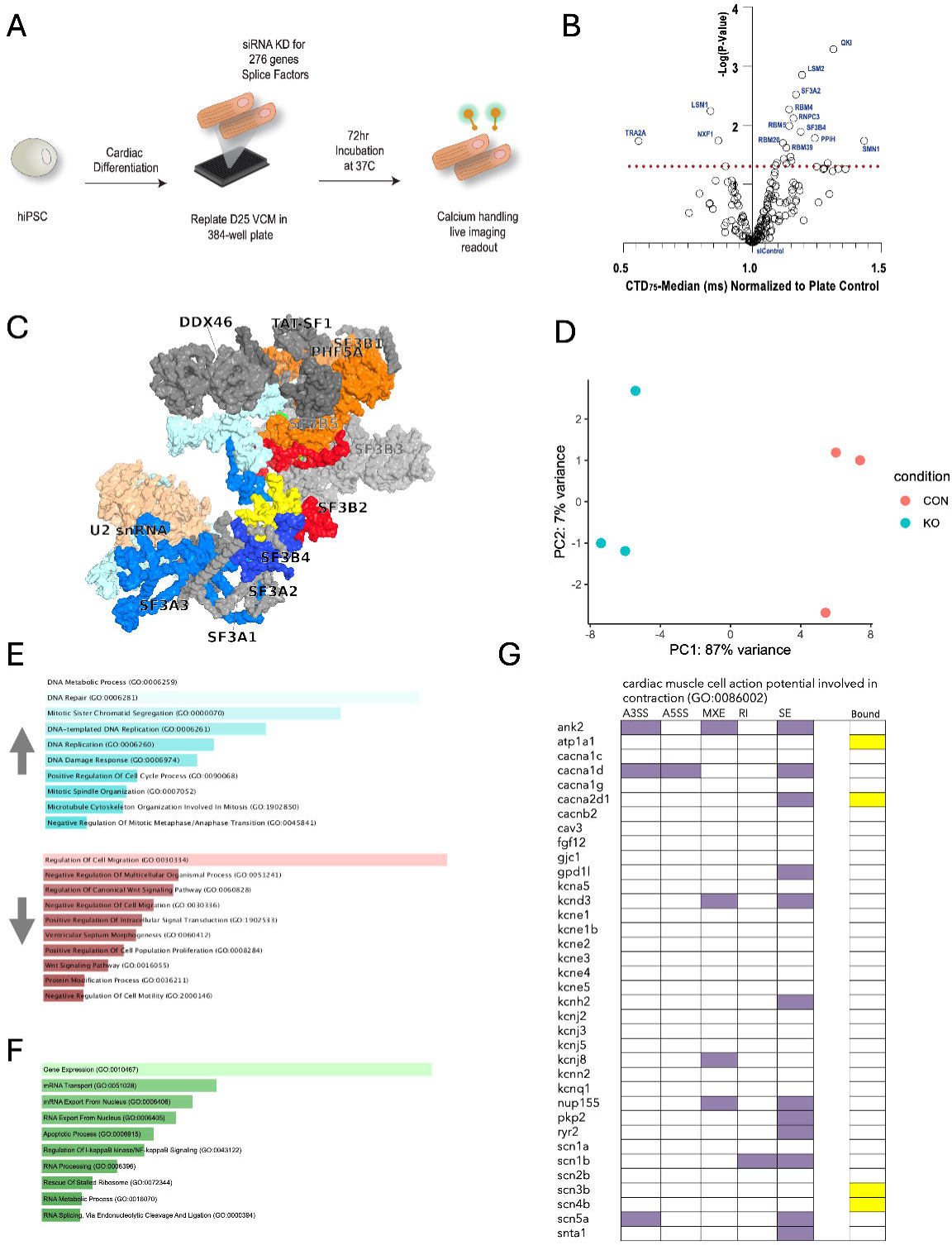
SF3B2 regulates cardiomyocyte calcium handling through alternative splicing of cardiac ion channel genes. **(A)** Schematic of the high-throughput splicing factor screen. Human iPSCs were differentiated to ventricular cardiomyocytes (VCMs), replated at day 25 into 384-well plates, transfected with siRNAs targeting 276 splicing factor genes, incubated for 72 hours, and assessed for calcium handling by live calcium imaging. Calcium transient duration at 75% repolarization (CTD75) was used as the functional readout. **(B)** Calcium transient duration screen results. Each point represents one splicing factor gene. x-axis: CTD75 median (ms) normalized to plate scrambled siRNA control; y-axis: –log_10_(*p*-value). Dashed red line indicates the significance threshold. U2 snRNP components (SF3A2, SF3B4) and other top hits (QKI, LSM2) are labeled. SF3B2 was previously identified as a screen hit regulating contractility [1]; its complex partners scored above the CTD75 significance threshold in this analysis. **(C)** Protein structure of the human 17S U2 snRNP complex (PDB: 6Y5Q) with screen hits highlighted in color [3]. Subunits of the SF3a subcomplex (SF3A2, SF3A3) and SF3b subcomplex (SF3B1, SF3B2, SF3B4, SF3B5) identified in the screen are indicated. **(D)** Principal component analysis (PCA) of bulk RNA-seq data from control (CON, red, *n* = 3) and SF3B2 knockdown (KD, green, *n* = 3) hiPSC-CMs. PC1 accounts for 87% of variance; PC2 accounts for 7%. **(E)** Gene ontology (GO) enrichment analysis of differentially expressed genes upon SF3B2 knockdown. Downregulated pathways (blue bars) include DNA metabolic process, DNA repair, mitotic sister chromatid segregation, DNA replication, cell cycle regulation, and spindle organization. Upregulated pathways (red bars) include regulation of cell migration, intracellular signal transduction, and ventricular septum morphogenesis. **(F)** GO enrichment analysis of transcripts with significantly retained introns upon SF3B2 knockdown (FDR < 0.01, inclusion level change > 0.1). Enriched terms include gene expression, mRNA transport, RNA export from nucleus, NF-kappaB signaling, RNA processing, rescue of stalled ribosome, and RNA splicing. **(G)** Heatmap of alternative splicing events and SF3B2 RNA binding across 36 genes annotated under the cardiac muscle cell action potential involved in contraction GO term (GO:0086002). This gene set includes ion channel genes as well as sarcomeric and structural genes (e.g., myosin heavy chains, troponins, tropomyosins, actinin). Columns represent five rMATS alternative splicing event types (A3SS, alternative 3’ splice site; A5SS, alternative 5’ splice site; MXE, mutually exclusive exons; RI, retained intron; SE, skipped exon) and RIP-seq binding enrichment (Bound). Colored cells indicate statistically significant events (rMATS: FDR < 0.05; RIP-seq: FDR < 0.05). *CACNA2D1* is highlighted as the convergent target exhibiting both skipped exon events and direct SF3B2 binding.

### SF3B2 Knockdown Disrupted Gene Expression and Alternative Splicing of Cardiac Contraction Genes

To elucidate the molecular consequences of SF3B2 loss, we performed deep bulk RNA-seq on SF3B2 knockdown and control hiPSC-CMs (n = 3 per group). SF3B2 knockdown efficiency was validated by qPCR, confirming 78% +/- 6% (mean +/- SD, n = 3) reduction of SF3B2 mRNA relative to scrambled siRNA controls. Principal component analysis (PCA) of the transcriptomic data revealed clear separation between control and SF3B2 knockdown groups, with PC1 accounting for 87% of the total variance (Fig. 1D). Differential expression analysis identified 2,683 genes with significantly altered expression (DESeq2, adjusted *p* < 0.05).

GO enrichment analysis separated the differentially expressed genes into coherent functional categories (Fig. 1E). Downregulated genes were enriched in cell cycle-related processes, including DNA metabolic process, DNA repair, mitotic sister chromatid segregation, DNA replication, DNA damage response, positive regulation of cell cycle, and mitotic spindle organization. Upregulated genes were enriched in regulation of cell migration, positive regulation of intracellular signal transduction, and ventricular septum morphogenesis. These expression changes are consistent with cell cycle arrest.

To determine whether SF3B2 loss disrupted pre-mRNA splicing, we performed rMATS analysis across all five alternative splicing event types. GO enrichment analysis of transcripts with significantly retained introns (FDR < 0.01, inclusion level change > 0.1) revealed enrichment in core RNA processing pathways, including gene expression, mRNA transport, RNA export from the nucleus, NF-kappaB signaling, RNA processing, rescue of stalled ribosome, and RNA splicing (Fig. 1F). The retained-intron transcripts thus converged on RNA processing and gene expression pathways.

To evaluate the impact of SF3B2 loss on cardiac-specific transcripts, we analyzed alternative splicing events within 36 genes annotated under the cardiac muscle cell action potential involved in contraction GO term (GO:0086002) across all five rMATS event types (Fig. 1G). Multiple cardiac ion channel genes exhibited significant splicing changes, including *CACNA1C*, *CACNA1D*, *CACNA2D1*, *CACNB2*, *KCNH2*, *KCNQ1*, *SCN5A*, and *RYR2*.

### Integration of RIP-seq and Splicing Data Identified CACNA2D1 as a Direct SF3B2 Target

We next asked which of these splicing changes reflected direct SF3B2 binding. RIP-seq was performed using an AAV6-delivered SF3B2-3xHA construct in hiPSC-CMs, with GFP-expressing cells as a control. This analysis identified 597 transcripts significantly enriched in the SF3B2-3xHA immunoprecipitation (DESeq2, FDR < 0.05). We integrated the RIP-seq binding data with the rMATS splicing analysis by including a “Bound” column in the cardiac action potential gene heatmap (Fig. 1G), enabling direct comparison of SF3B2 binding status with alternative splicing events for each gene.

*CACNA2D1* was the only gene in this set that was both directly bound by SF3B2 and alternatively spliced upon knockdown (Fig. 1G). *CACNA2D1* encodes the alpha-2-delta-1 auxiliary subunit of voltage-gated calcium channels, which regulates channel trafficking to the plasma membrane and modulates calcium current density [22, 23]. Mutations in *CACNA2D1* have been associated with cardiac arrhythmias and short QT syndrome [23]. SF3B2-dependent missplicing of *CACNA2D1* is therefore a candidate mechanism for the calcium handling phenotype, though functional validation of the individual isoforms remains to be performed.

## Discussion

SF3B2 knockdown altered calcium transient duration in hiPSC-CMs, and integration of RNA-seq, splicing analysis, and RIP-seq converged on *CACNA2D1* as a direct SF3B2 target whose missplicing may underlie this phenotype.

Unlike the accessory RBPs studied to date—RBM20 [11], MBNL1 [3], RBFOX2 [12], QKI [14]—SF3B2 is a core spliceosome component, raising the question of how it exerts transcript-specific effects. Precedent for this comes from studies of SF3B1, a paralog of SF3B2 within the SF3b subcomplex, whose somatic mutations in myelodysplastic syndromes cause aberrant 3’ splice site selection in a transcript-specific manner [24, 25]. *CACNA2D1* is an established regulator of L-type calcium channel trafficking and surface expression [22], and variants in the *CACNA2D1* locus have been associated with cardiac arrhythmias [23], lending biological plausibility to its identification as a downstream effector of SF3B2.

One explanation is that transcripts with weak or degenerate branch point sequences are disproportionately sensitive to reduced SF3B2 levels, given the SF3b subcomplex’s role in branch point recognition during early spliceosome assembly [17]. Such transcripts would be prone to exon skipping, intron retention, or alternative splice site usage even when most splicing events proceed normally. The identification of six U2 snRNP subunits in the screen reinforces this interpretation: reduced U2 complex integrity, rather than loss of any single subunit, appears to underlie the calcium handling phenotype. The concurrent cell cycle arrest observed upon SF3B2 knockdown may reflect the broader requirement for intact spliceosomal function in proliferation-associated transcripts, as has been reported in other cellular contexts [26]. Because hiPSC-CMs retain proliferative capacity, cell cycle arrest could independently alter calcium handling properties through changes in ion channel expression or cellular maturation state. The 72-hour timepoint used in the functional screen and the 48-hour timepoint used for RNA-seq may also differ in the degree of cell cycle disruption. It therefore remains possible that some portion of the calcium transient phenotype reflects secondary consequences of cell cycle arrest rather than direct effects of ion channel missplicing, and future experiments that uncouple these two effects will be informative.

Beyond calcium handling, our findings position SF3B2 within the broader splicing circuitry that governs cardiomyocyte maturation. Postnatal cardiac maturation is driven in part by organ-wide alternative splicing transitions [5], and dedicated splicing regulators such as RBFox1 control this process [10]; the immature, fetal-like state of hiPSC-CMs is itself attributed in part to failed execution of the perinatal maturation program [7, 8], of which alternative splicing is an integral component [5, 10]. Although the cell cycle arrest that follows SF3B2 knockdown may be a nonspecific consequence of broad spliceosomal disruption rather than a physiological maturation signal, both cell cycle exit and switches in ion-channel and calcium-handling splicing are hallmarks of the fetal-to-adult transition, raising the speculative possibility that SF3B2 levels contribute to coupling cell cycle exit with acquisition of a mature contractile phenotype. Because maturation state and proliferative or regenerative capacity are inversely related in cardiomyocytes, clarifying how core spliceosome components such as SF3B2 tune these splicing programs may inform strategies both to enhance hiPSC-CM maturation for disease modeling and to modulate regenerative potential in the injured heart.

## Future Directions

Several priorities for future work follow directly from the present study’s limitations. Because our experiments used relatively immature hiPSC-CMs [27], an important next step is to test whether the SF3B2-dependent splicing changes and the *CACNA2D1* phenotype persist in more mature or adult cardiomyocytes, whose splicing landscape may differ. As siRNA-mediated knockdown is transient, stable or genetic loss-of-function models will be needed to define the consequences of sustained SF3B2 depletion. Direct targets should be re-mapped at endogenous SF3B2 levels—for example by eCLIP—using more stringent controls than the GFP comparison employed here, such as an epitope-tagged unrelated nuclear protein, to exclude non-physiological binding arising from CMV-driven overexpression. Establishing causality will further require functional validation of the individual *CACNA2D1* splicing isoforms produced upon SF3B2 knockdown and their specific effects on calcium-channel trafficking and calcium handling, together with confirmation of the key findings across additional iPSC lines and genetic backgrounds. Several of these goals are addressed by the experiments outlined below.

Three-dimensional macroscale cardiac tissue or organoid constructs generated would allow calcium hanlding and contractility assessment at more physiological relevant level [28, 29, 30]. A cardiomyocyte-specific *Sf3b2* conditional knockout mouse has been generated and cardiac phenotyping is underway. Isoform-specific rescue experiments will test whether individual *CACNA2D1* splice variants are sufficient to restore calcium handling in SF3B2-depleted hiPSC-CMs.

## Acknowledgments

This work was supported by funding from the American Heart Association (23TPA1058685) and the National Institutes of Health (R01HL171205).

## Author Contributions

C.K., S.M., H.W. conceived and design the study. S.M., N.Z., M.K., performed the experiments. H.W., D.L., C.K., wrote the manuscript. All authors reviewed and approved the final version.

## Data Availability

All analysis code is available at https://github.com/kwon-lab/SF3B2-splicing.

